# Disrupted Post-Transcriptional Regulation of Gene Expression as A Hallmark of Fatty Liver Progression

**DOI:** 10.1101/2024.09.02.610906

**Authors:** Shohei Takaoka, Marcos E Jaso-Vera, Xiangbo Ruan

**Affiliations:** Division of Endocrinology, Diabetes and Metabolism, Johns Hopkins University School of Medicine, Baltimore, MD, USA; Institute for Fundamental Biomedical Research, Johns Hopkins All Children’s Hospital, St. Petersburg, FL, USA

## Abstract

It is known that both transcriptional and post-transcriptional mechanisms control the messenger RNA (mRNA) levels. Compared to transcriptional regulations, our understanding of how post-transcriptional regulations adapt during fatty liver progression at the whole transcriptome level is unclear. While traditional RNA-seq analysis uses only reads mapped to exons to determine gene expression, recent studies support that intron-mapped reads can be reliably used to estimate gene transcription. In this study, we analyzed differential gene expression at both exon and intron levels using two liver RNA-seq datasets from mice that were fed a high-fat diet for seven weeks (mild fatty liver) or thirty weeks (severe fatty liver). We found the correlation between gene transcription and mature mRNA levels was much lower in mice with mild fatty liver as compared with mice with severe fatty liver. This result indicates broad post-transcriptional regulations for early fatty liver and such regulations are comprised for severe fatty liver. Specifically, Gene Ontology analysis revealed that genes involved in synapse organization and cell adhesion were transcriptionally upregulated, while their mature mRNAs were unaffected in the mild fatty liver. Further characterization of post-transcriptionally suppressed genes at early fatty liver revealed that their mRNAs harbor significantly longer 3’ UTR, one of the major features that may subject RNA transcripts to non-sense mediated RNA decay (NMD). We further show that the expression of representative genes that were post-transcriptionally suppressed were upregulated in mice with hepatocyte-specific defect of NMD. Finally, we provide data supporting a time-dependent decrease in NMD activity in the liver of a diet-induced metabolic dysfunction-associated fatty liver disease mouse model. In summary, our study supports that NMD is essential in preventing unwanted/harmful gene expression at the early stage of fatty liver and such a mechanism is lost due to decreased NMD activity in mice with severe fatty liver.

## INTRODUCTION

Overnutrition-associated obesity increases all forms of cardiometabolic diseases, including hyperlipidemia, diabetes and fatty liver diseases. In response to overnutrition, tissues/organs adapt gene expression/activity to maintain metabolic homeostasis. With the overnutrition-associated stress continuing, tissues/organs exhaust the capacity to adapt gene expression/activity to maintain homeostasis, leading to metabolic diseases. In the past decades, tremendous progress has been made in understanding how genes are transcriptionally regulated in response to major nutrients/hormones. For example, transcription factor SREBP1c controls feeding associated de novo lipogenesis^1^, and SREBP2 senses cellular cholesterol content to promote cholesterol synthesis^2^, and PPAR-α activation promotes fatty acid catabolism^3^. However, at the RNA level, gene expressions are controlled by both transcriptional and post-transcriptional mechanisms. These post-transcriptional mechanisms include, splicing and processing of pre-mRNA, transportation of mRNA and nuclear export, translation of mRNA and ultimately, the degradation of mRNA. Compared to well-studied transcriptional regulation, our understanding of post-transcriptional mechanisms during diet-induced obesity is much less clear.

The importance of post-transcriptional mechanisms in metabolic diseases are emerging. Recent mouse genetic studies found that knocking out of key components of RNA decay machinery and some RNA-binding proteins (RBPs), and splicing factors, resulted in obesity, abnormal adipose tissue metabolism and liver diseases^4-7^. Despite these progresses, how post-transcriptional mechanisms adapt to overnutrition is largely unclear, especially at the whole transcriptome level due to technical limitations. While ChIP-seq and RNA-seq have been the standard procedure to study transcriptional regulations at tissue/organ levels, most of the established methods for chasing RNA-decay in cultured cells, like blocking transcription by actinomycin D, or BrU labeling of nascent RNA, are not practical or with limitations in animals^8^.

The recent advances in high-throughput RNA sequencing have provided huge advantages in defining differential gene expressions at multiple levels. In addition to determining total gene expression levels, extensive RNA-seq analysis can also determine splicing patterns and alternative usage of 3 prime and 5 prime ends. One unique feature of RNA-seq data is that they contain reads that can be mapped to both introns and exons, especially when the RNA-seq library was prepared using total RNA. As such, reads mapped to introns can serve as an estimate of transcription activity, and reads mapped to exons can serve as expression of mature mRNA. When analyzing RNA-seq data, by comparing the differential gene expressions at the intron level with those at the exon level, the effects of post-transcriptional regulations can be measured. This strategy was originally proposed by Gaidatzis et al^9^. and has been applied to determine the role of post-transcriptional regulations in many pathophysiological conditions.

In this study, we established a robust RNA-seq pipeline that determines gene expression at both the intron and exon levels and applied the pipeline to analyze published RNA-seq datasets representing hepatic gene expressions in mice with mild and severe fatty liver^10,11^. Our results revealed broad post-transcriptional regulations in mild fatty liver, and these post-transcriptional regulations were lost in server fatty liver. We further provide evidence that non-sense mediated RNA decay (NMD) is an important mechanism contributing to post-transcriptional regulations in mild fatty liver.

## RESULTS

### Broad post-transcriptional gene suppression at early stage of fatty liver

To systemically define the contribution of post-transcriptional regulations in fatty liver-associated gene expression changes, we searched for published RNA-seq datasets that represent liver gene expression at different stages of fatty liver. Two RNA-seq datasets deposited in NCBI were used: GSE88818 contains RNA-seq results from the liver of mice fed with either a normal chow diet or a high-fat diet (HFD) for seven weeks representing the stage of mild fatty liver^10^; GSE121340 contains RNA-seq results from the liver of mice fed with either a normal chow diet or an HFD for thirty weeks, thus represent severe fatty liver^11^. The RNA-seq libraries for both datasets were prepared using total RNA (ribosomal RNA depleted), with total reads ranging from 50-90 M. To determine differentially expressed genes (DEG) at both intron and exon levels, we established the RNA-seq pipeline as shown in (**Figure 1)**. Compared with default RNA-seq analysis, in which RNA-seq reads were only mapped to exon regions, our pipeline map reads to either exon or introns. As there are usually multiple RNA transcripts for a single gene, we only use reads that always map to exons (constitutive exons) or always map to introns (constitutive introns) based on the Ensembl annotation (**see Methods**). We detected a total of 397 exon level, and 595 intron level DEGs for mild fatty liver, and a total of 1288 exon level, and 993 intron level DEGs for severe fatty liver (**Supplementary Table 1**). To globally define the contribution of post-transcriptional regulations, we performed a correlation analysis to determine how intron-level gene expression changes (Δintron) correlate with exon-level gene expression changes (Δexon) (**see Methods**). As shown in **Figure 2A and 2B**, the R^2^ for this correlation is 0.56 for mild fatty liver and 0.83 for severe fatty liver. This result suggests that gene expression changes at the intron level were less efficiently transferred to gene expression changes at the exon level at early stage of fatty liver, as compared to what happened in severe fatty liver. It also indicates that broad post-transcriptional regulations at early stage of fatty liver, and gene expressions at severe fatty liver are mainly determined by gene transcription.

**Figure 1.**
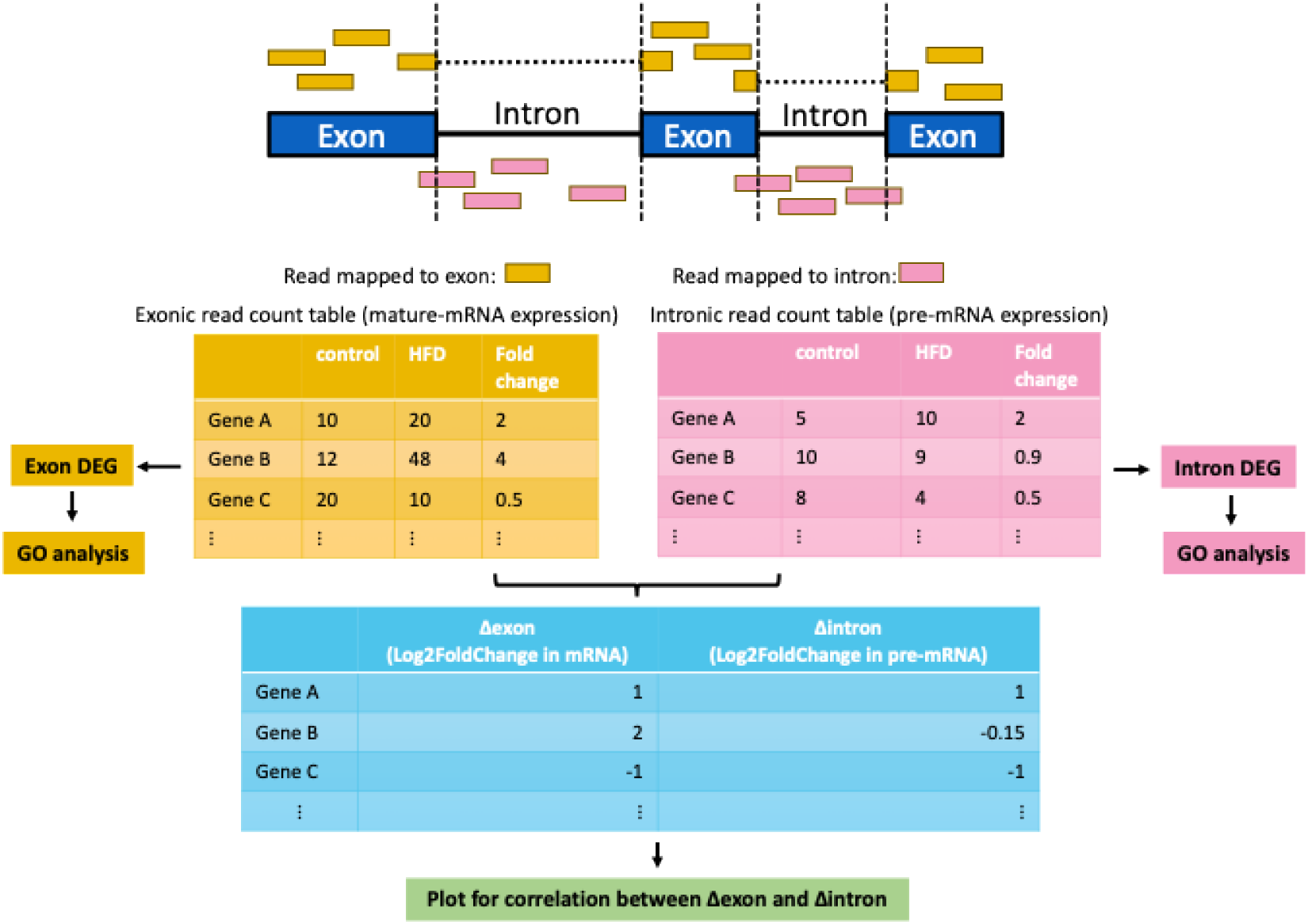
Schematics of the analysis workflow. RNA-seq reads are mapped to the genome, and those reads assigned to exon and intron regions are aggregated to calculate exon and intron counts for each gene, respectively. The obtained count table is used for differential expression analysis by DESeq2 to calculate Log 2Fold change in mature mRNA (Δexon) and Log2FoldChange in pre-mRNA (Δintron). Scatter plots of Δexon and Δintron are drawn for each data set to evaluate how much the change in transcription (Δintron) contributes to the variation in expression in mature mRNA seen in each data set.

**Figure 2.**
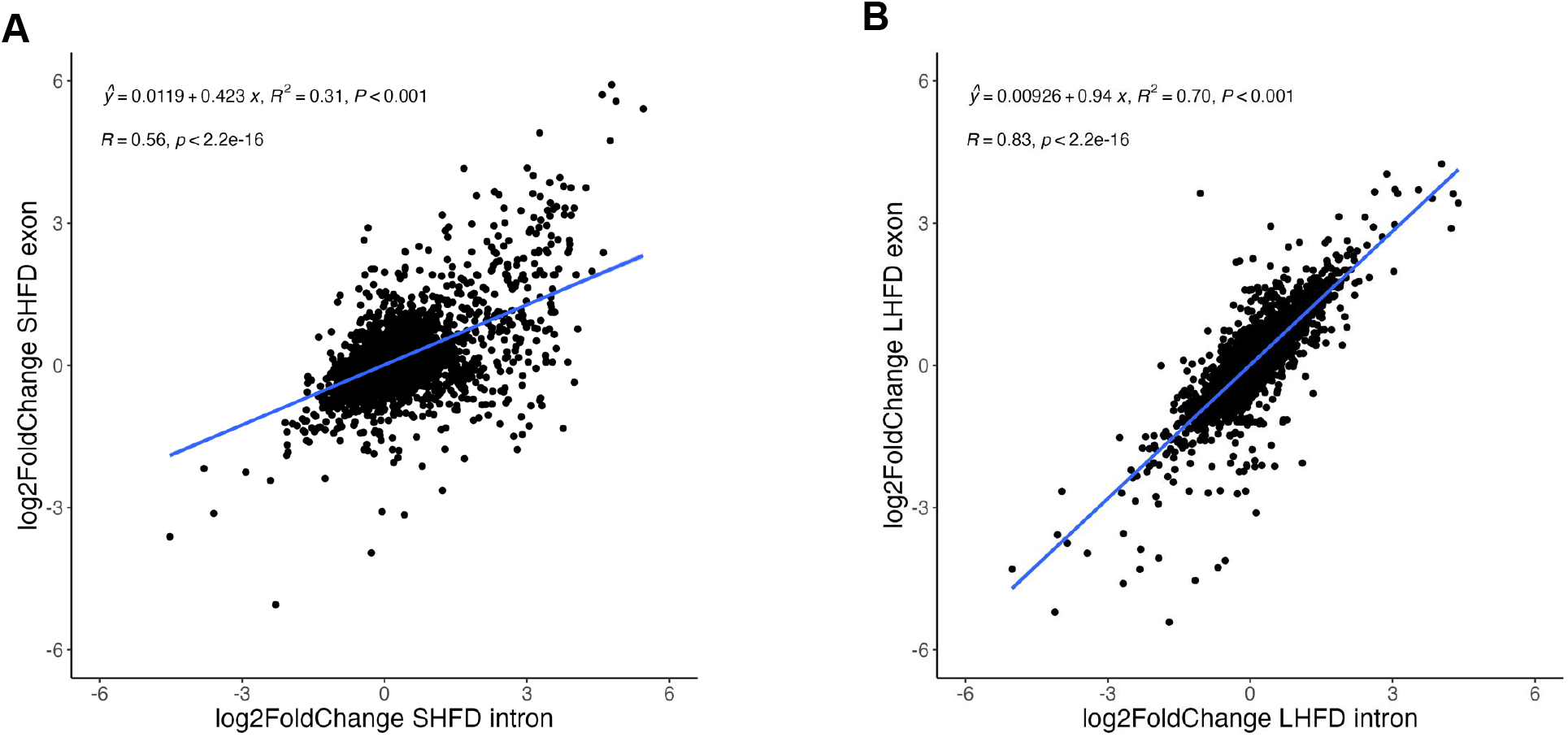
**A**. The correlations between Δexon and Δintron for DEG of short-term HFD (SHFD). **B**. The correlations between Δexon and Δintron for DEG of Long-term HFD (LHFD). Scatter plots of the results of analyzing each dataset with the analysis pipeline shown in Figure 1.

### Genes involved in synapse organization and cell adhesion were post-transcriptionally suppressed at mild fatty liver

To define the major genes/pathways that undergo post-transcriptional regulations at mild fatty liver, we performed GO term analysis for both our intron and exon DEG datasets. As shown in **Figure 3**, genes with exon-level upregulation were enriched in the fatty acid metabolic process and organic acid biosynthetic process (**Figure 3A**), and genes with exon-level downregulation were mainly involved in fatty acid catabolism and xenobiotic metabolic process at mild fatty liver (**Figure 3B**), which is consistent with our current knowledge about how mouse livers response to HFD at early stage^10,12^. While GO term analyses for genes with intron-level downregulation (**Figure 3D**) were similar to GO terms for genes with exon-level downregulation (**Figure 3B**), we found that GO terms for genes with intron level upregulation were represented by synapse organization and cell adhesion (**Figure 3C**), which is not observed in top GO terms for genes with exon-level upregulation at mild fatty liver (**Figure 3A**). These results are consistent with our correlation analysis as shown in **Figure 2A**. It also suggests genes involved in synapse organization and cell adhesion were post-transcriptionally suppressed, and this is the major factor contributing to the lower correlation between gene transcription and mature mRNA levels at early stage of fatty liver. We next performed the same GO term analysis in severe fatty liver. We found that at both intron and exon level, upregulated genes were highly enriched for GO terms related to immune activation, and downregulated genes were highly enriched for GO terms related to small molecule catabolic process (**Figure 4 A-D**). These results are in line with our correlation analysis as shown in **Figure 2B** and are also consistent with our knowledge that long-term HFD induces dramatic inflammatory responses and suppressed fatty acid catabolism and drug metabolism^11^. Taken together, our GO term analyses are consistent with our correlation analysis. It also revealed that genes involved in synapse organization and cell adhesion were post-transcriptionally suppressed at the early stage of fatty liver.

**Figure 3.**
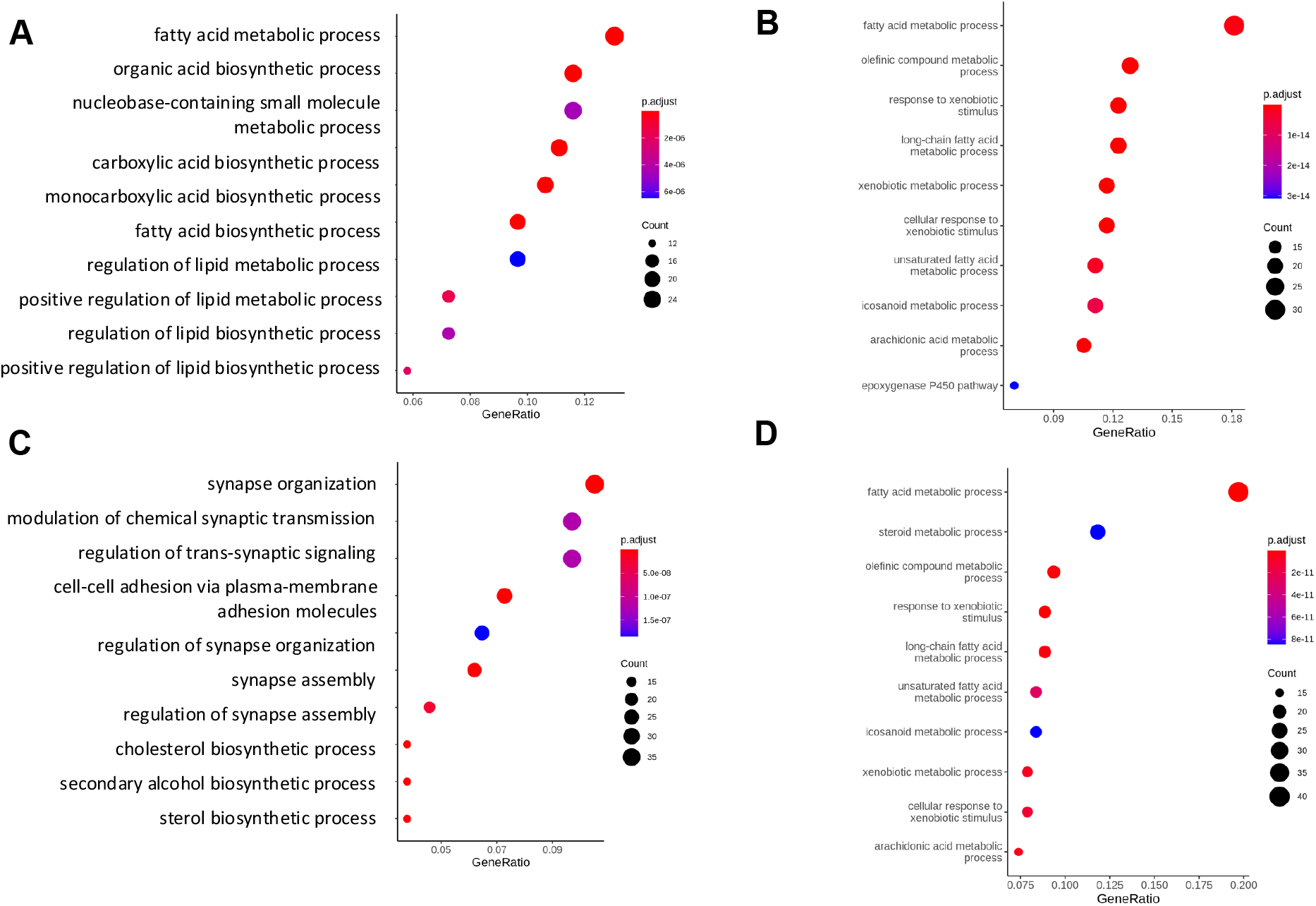
**A**. GO term analysis for exon-level upregulated genes of short-term HFD, **B**. GO term analysis for exon-level downregulated genes of short-term HFD, **C**. GO term analysis for intron-level upregulated genes of short-term HFD, **D**. GO term analysis for exon-level downregulated genes of short-term HFD.

**Figure 4.**
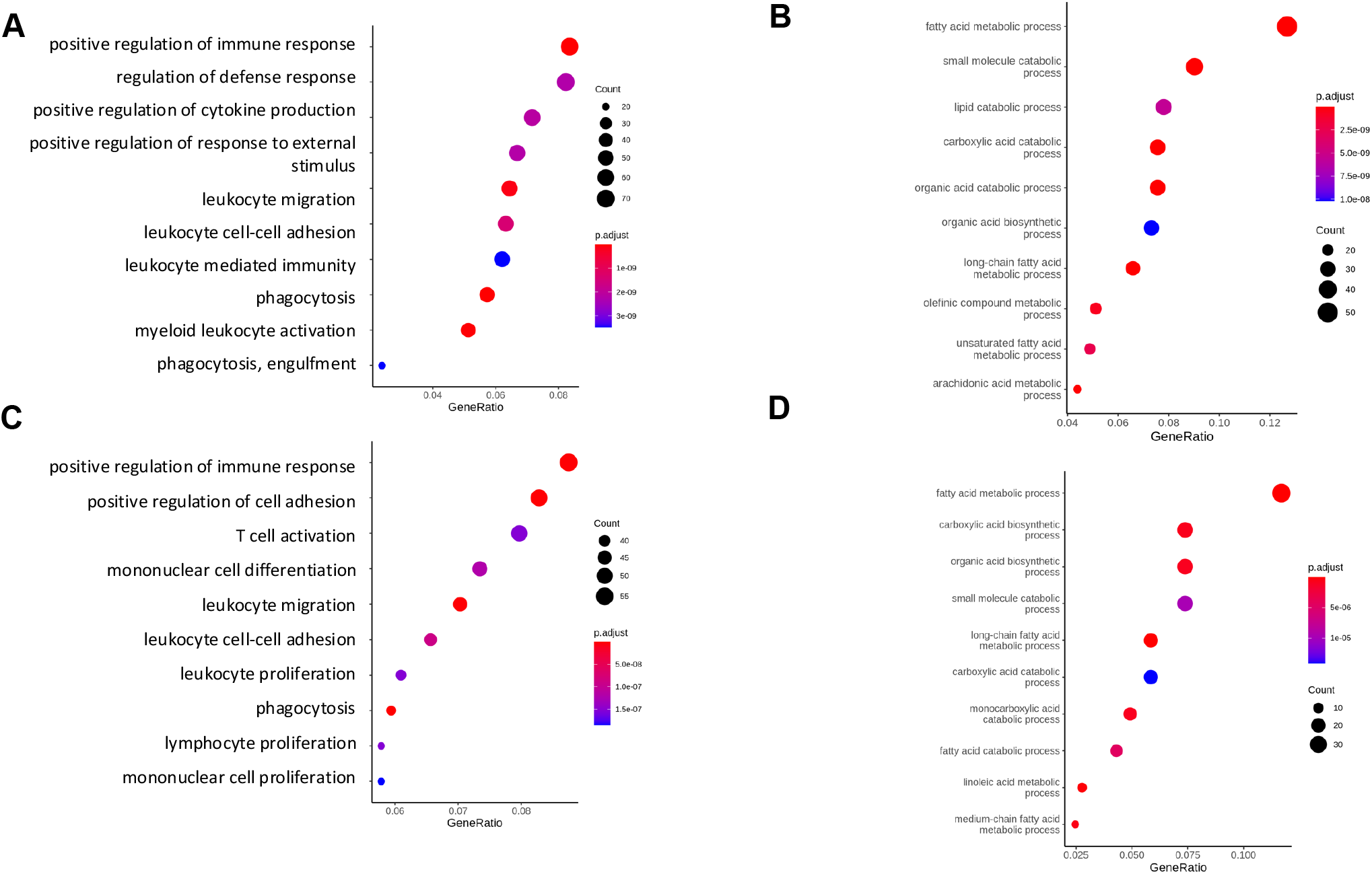
**A**. GO term analysis for exon-level upregulated genes of long-term HFD, **B**. GO term analysis for exon-level downregulated genes of long-term HFD, **C**. GO term analysis for intron-level upregulated genes of long-term HFD, **D**. GO term analysis for exon-level downregulated genes of long-term HFD.

### Post-transcriptionally suppressed genes at mild fatty liver have long UTRs

To explore the potential mechanism underlying post-transcriptional regulations during fatty liver progression, we next asked if differentially expressed genes at early obesity shared any common features that may subject them to post-transcriptional regulations. Given the significance of the untranslated region (UTR) of mRNA in post-transcriptional regulations, we first systemically determined the 3’ UTR length distribution among the intron/exon level DEGs at both short-term and long-term HFD setting. As shown in **Figure 5A**, we found that only genes with intron-level upregulation at short-term HFD showed a significantly longer 3’ UTR when compared with the average UTR length of all liver expressed genes, or DEGs at other conditions. This result suggests that the long 3’ UTR may explain post-transcriptional suppression of many intron-level upregulated genes at short-term HFD. We also found that genes with intron/exon-level downregulation at either short or long-term HFD showed shorter 3’ UTR when compared with the average UTR length of all liver expressed genes (**Figure 5**), suggesting these genes may be subject to lower level of post-transcriptional regulation. This is in line with our observation that Go terms for downregulated intron/exon DEGs at short and long-term HFD were similar. We further determined the 5’ UTR distribution in the same setting (**Figure 6**) and found that only genes with intron-level upregulation at short-term HFD showed a significantly longer 5’ UTR when compared with the average UTR length of all liver expressed genes (**Figure 6A**).

**Figure 5.**
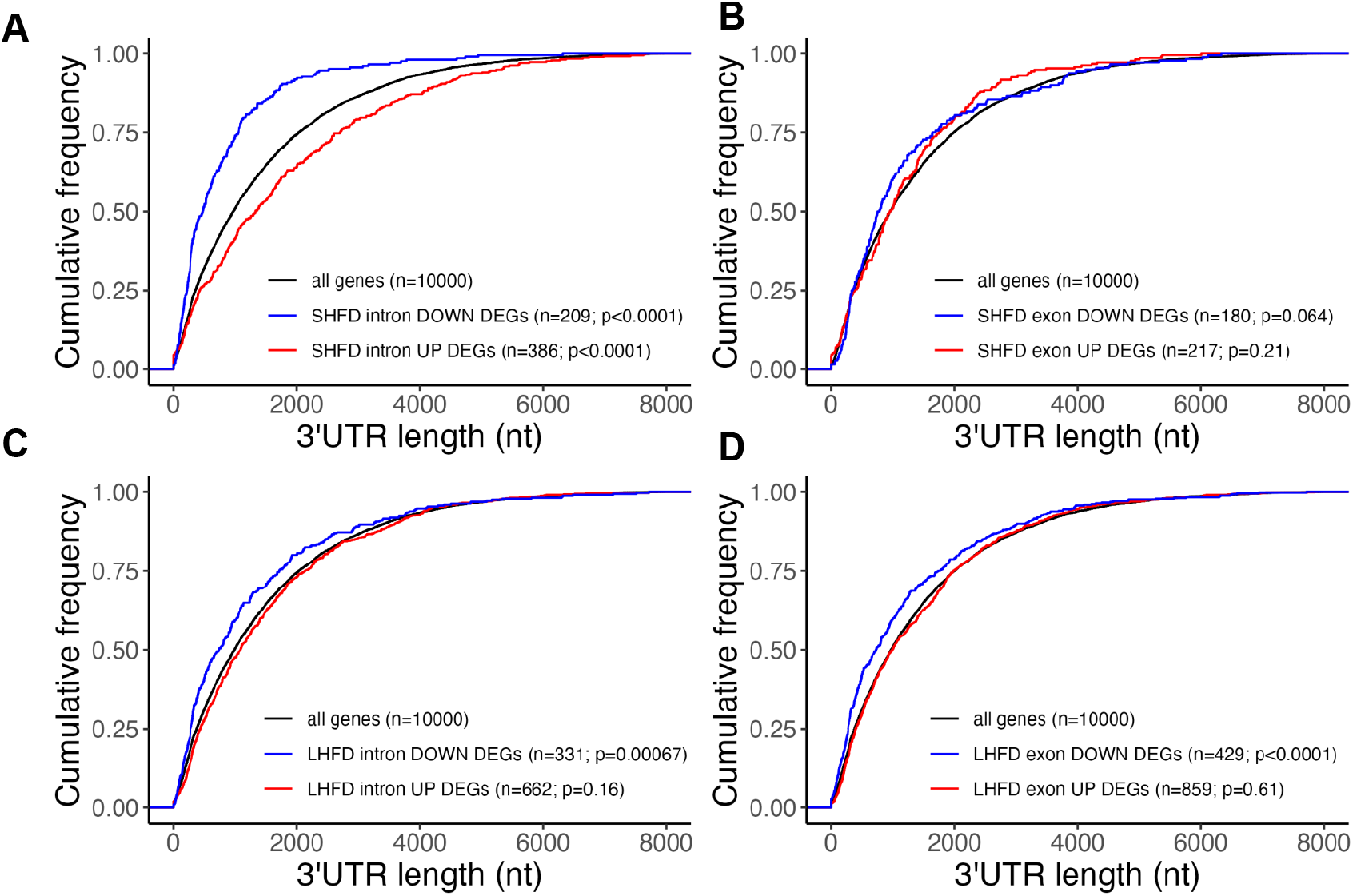
**A**. 3’ UTR distribution for intron-level DEGs of short-term HFD. **B**. 3’ UTR distribution for exon-level DEGs of short-term HFD. **C**. 3’ UTR distribution for intron-level DEGs of long-term HFD. **D**. 3’ UTR distribution for exon-level DEGs of long-term HFD.

**Figure 6.**
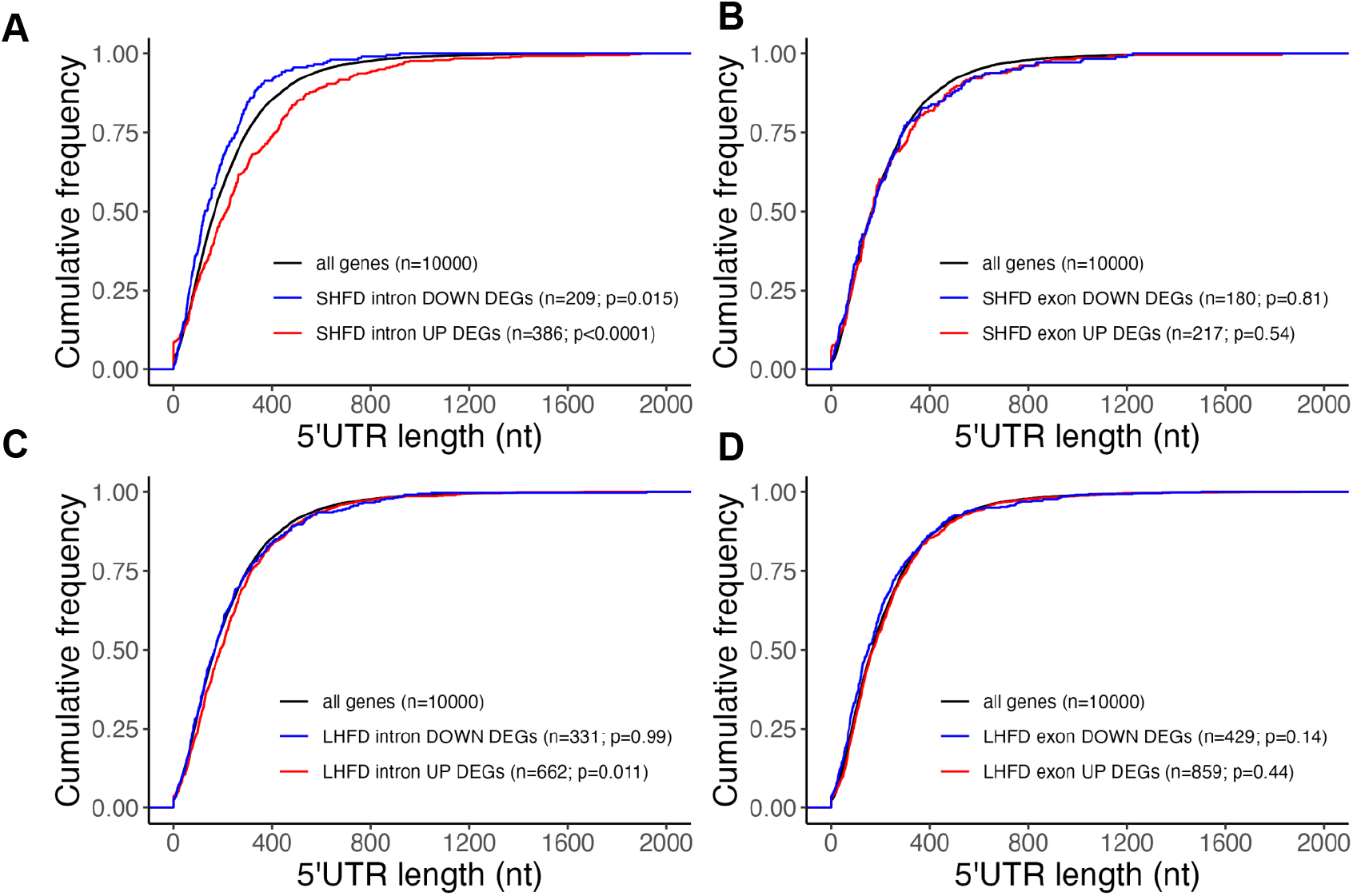
**A**. 5’ UTR distribution for intron-level DEGs of short-term HFD. **B**. 5’ UTR distribution for exon-level DEGs of short-term HFD. **C**. 5’ UTR distribution for intron-level DEGs of long-term HFD. **D**. 5’ UTR distribution for exon-level DEGs of long-term HFD.

### Post-transcriptionally suppressed genes in early fatty liver are NMD targets

Having identified long 3’ UTR as a major feature of post-transcriptionally suppressed genes at early fatty liver, we next search for post-transcriptional mechanisms that were reported to correlate with 3’ UTR length. We noticed that non-sense mediated RNA decay (NMD), an RNA quality control mechanism not only targets mRNA with pre-mature stop code but also shows a strong preference to degrade RNA transcripts with long 3’ UTRs^13^. To test if post-transcriptionally suppressed genes at early fatty liver are subjected to NMD, we prepared mice with hepatocyte-specific knockout of Upf2, a core component of NMD, by applying adeno-associated virus expressing Cre recombinase driven by thyroxine binding globulin (TBG) promoter (AAV8-TBG-Cre) in Upf2^flox/flox^ mice^14^. As shown in **Figure 7A**, three weeks after AAV8-TBG-Cre injection, this strategy successfully depleted Upf2 protein expression in the liver. We next determined the expression of several representative genes, including Robo1, Gabrb3, Gpc6, Ephb2 and Dlg2, based on our analyses that: (1) their expressions were post-transcriptionally suppressed at short-term HFD, and upregulated at both intron and exon level in long-term HFD; (2) their involvement in synapse organization and cell adhesion process based on our GO term analysis and (3) their extreme long 3’ UTRs. As shown in **Figure 7B**, we found that Upf2 knockout resulted in strong upregulation of Robo1, Ephb2, and Dlg2, suggesting they may be subjected to NMD. Given that the broad post-transcriptional regulations were mainly observed at early stage of fatty liver, but not at severe fatty liver, we next asked if hepatic NMD activity is compromised during fatty liver progression. We thus measured the phosphorylation of Upf1, which is commonly used as an estimate of NMD activity^15^, in the liver of mice fed with Gubra-Amylin NASH (GAN) diet^16^ for 3 months or 6 months. As shown in **Figure 7C**, we found Upf1 phosphorylation was gradually suppressed upon GAN diet feeding with a substantial decrease at 6 months, when NASH-like phenotypes developed in these mice.

**Figure 7.**
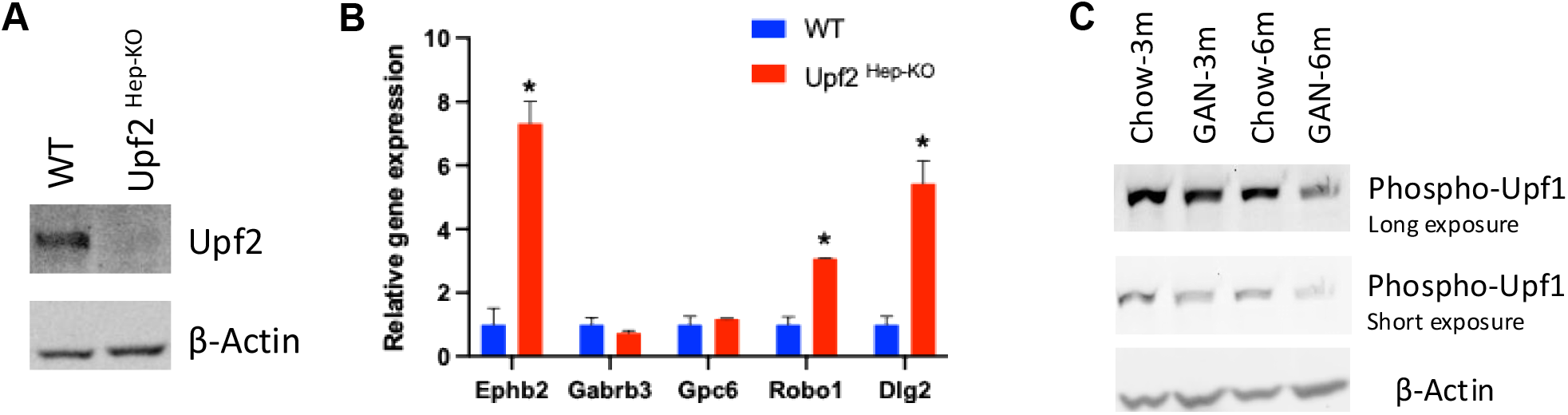
**A**. Western blot analysis detecting Upf2 and β-Actin in the liver of WT or Upf2 ^Hep-KO^ mice. **B**. qPCR detecting the relative expression of genes indicated (n=4 for WT, and n=2 for Upf2 ^Hep-KO^. Data shown as the mean ± SEM, *p<0.05. **C**. Western blot analysis detecting phospho-Upf1 (Ser1127) or β-Actin in mice fed with chow or GAN diet for 3 or 6 months. Each lane represents samples pooled from 5-7 mice.

## DISCUSSION

Mammalian cells utilize multi-level regulatory mechanisms to control gene expression. At the mRNA level, gene expression is controlled by both transcription and degradation. In this study, we employed an RNA-seq data analysis pipeline that determines gene expression at both intron and exon levels, to study how gene expression in the liver adapts to high-fat diet feeding. Our major finding includes: (1) in the early stage of HFD-induced fatty liver, there are broad post-transcriptional regulations to suppress genes involved in cell adhesion and synapse organization; (2) at the late stage of HFD-induced fatty liver, gene expression is mainly controlled by transcription; (3) post-transcriptionally suppressed genes at early fatty liver have extreme long 3 primer UTRs and other features including longer 5 primer UTR and longer introns, making them more susceptible to post-transcriptional regulation; (4) we provide evidence that post-transcriptionally suppressed genes at early fatty liver are subject to NMD, and suppressed NMD activity may partially explain loss of post-transcriptional regulations in severe fatty liver.

It is well known that upon stress/stimulations, cells adapt by controlling gene expression/activity at multiple layers. Our study here revealed upon HFD feeding, mice utilize both transcriptional and post-transcriptional mechanisms to maintain metabolic homeostasis. Specifically, we found that genes enriched for synapse organization and cell adhesion were post-transcriptionally suppressed at early stage of fatty liver. These genes, including Robo1, Gabrb3, Gpc6, Ephb2 and Dlg2, usually do not express in healthy liver, were reported to be upregulated in metabolic dysfunction-associated steatohepatitis or liver fibrosis. Especially, increased Robo1 and Ephb2 expression has been experimentally proved to promote MASH and liver fibrosis^17,18^. Our results suggest that post-transcriptional regulations represent a checkpoint to suppress the expression of unwanted/harmful genes in the liver upon HFD-induced nutritional stress. In our setting, short term HFD feeding (7-8 weeks) very likely represents a turn point, after which metabolic disease phenotypes may develop. Indeed, it is well accepted that while with increased body weight and mild fatty liver, mice fed with HFD for 8 weeks show no significant metabolic defects^19^.

The finding that post-transcriptionally regulated genes upon HFD feeding in the liver show features including long 3’ UTR, suggests they may be subject to NMD regulation. Given many of these genes are induced in disease status and promote liver disease progression, their expression is thus under tight surveillance by mechanism such as NMD. Indeed, for genes that are mainly controlled by transcriptional regulations, such genes with exon-level downregulation at the early stage of fatty liver, have significantly shorter 3’ UTR as compared with all liver expressed genes. These findings suggest that 3’ UTR length alone may determine to what extent a gene’s expression may be subject to post-transcriptional regulations such as NMD. Our result that Upf1 phosphorylation was maintained at early stage of fatty liver but lost at severe fatty liver, supports that loss of NMD activity may explain low post-transcriptional regulations at severe fatty liver diseases and possibly other chronic liver diseases including MASH, liver fibrosis, and liver cancer. It was reported that NMD activity can be suppressed by many environmental factors, including ER stress, a known feature of diet-induced fatty liver^20^. Our observation also suggests that restoring NMD activity may show a protective effect in liver diseases by suppressing the expression of disease-related genes, such as Robo1 and Ephb2.

## MATERIAL & METHOD

### Reanalysis of publicly available RNA-seq data

We downloaded 6 samples from dataset GSE88818, obtained from 19-week-old control mice (n = 3) fed a chow diet and 19-week-old obese mice (n = 3) fed HFD (D12327, Research Diet, 40% fat by calories). HFD feeding was started at 12 weeks of age and continued for 7 weeks. We also downloaded 6 samples from dataset GSE121340, obtained from 36-week-old control mice (n = 3) fed a chow diet and 36-week-old obese mice (n = 3) fed HFD (D12492, Research Diet, 60% fat by calories). HFD feeding was started at 6-7 weeks of age and continued for 29-30 weeks. The Fastq files were downloaded and prepared using the SRA Toolkit (version 2.11.1) [https://www.ncbi.nlm.nih.gov/books/NBK569238/]. Briefly, the prefetch command was used to download the SRA files of the dataset and convert them into FASTQ format using the fasterq-dump command. The quality of the Fastq files was evaluated using FastQC (version 0.11.8) [http://www.bioinformatics.babraham.ac.uk/projects/fastqc/], and adapters and low-quality reads were trimmed or removed using Trimmomatic (version 0.39) [PMID: 24695404]. The filtered reads were then mapped to the GRCm 39 Ensembl mouse genome using STAR (version 2.7.8a)^21^. Annotation files for gene regions to obtain mature mRNA and pre-mRNA count data were created as follows. From the annotation file of mouse genome, transcripts whose transcript source was ensembl or ensembl_havana were extracted. From there, exon regions common to all transcripts for each gene were extracted and designated as constitutive exon regions. For the creation of the intron region definition file, the region between adjacent exons was extracted for each gene. Gene-level counts for mature mRNA and pre-mRNA were generated using featureCounts from Subread (version 2.0.0)^22^ by using uniquely mapped reads in the union exon region or the union intron region, respectively. In each dataset, genes with two or fewer samples with raw counts greater than one were pre-filtered out. The pre-filtered raw count matrix was normalized by DESeq2 (version 1.36.0)^23^ on R (version 4.1.0). Differentially expressed genes were defined as genes with false discovery rate (FDR) <= 0.05 and |Log2FC| >=0.58. DEGs with top 10,000 in baseMean calculated by DESeq2 were used as an input for gene ontology (GO) enrichment analysis using cluster Profiler (version 4.0.5)^24^. In GO enrichment analysis, GO terms with adjusted p-value <= 0.01 were considered significant.

### Mouse study

Upf2^flox/flox^ mice were described as previously^25^. AAV8-TBG-Cre and AAV8-CMV-GFP control were purchased from Vector Biolabs. Six weeks male Upf2^flox/flox^ mice were injected with 2 × 10^11^ genome copies/mouse. Three weeks after AAV injection, liver tissues were harvested for RNA and protein extraction. For GAN diet study, 8 weeks C57BL/6J mice were fed with normal chow or GAN diet (Research Diets, D09100310) for 3 months or 6 months, then the liver tissue were harvested for western blot analysis. Mouse experiments were approved by the Johns Hopkins University Animal Care and Use Committee (ACUC).

### RNA extraction and qPCR

Total RNA was extracted and purified using the Qiagen RNeasy Mini Kit (Qiagen Cat# 74,106) following the manufacturer’s instructions and performing the on-column DNA digestion. 200 -500 ng of total RNA was used for cDNA synthesis with the SuperScript III First-Strand Synthesis SuperMix for qRT-PCR (Invitrogen Cat# 11,752,050). cDNA was diluted 3-5-fold with RNase-free water. qPCR reactions consisted of 7.5 µl of the Power SYBR Green PCR Master Mix (ThermoFisher Cat# 4,368,702), 1.5 μL of cDNA, and 6 µl of 0.5 µM primers (primers are listed in Table X). qPCR was performed using a ThermoFisher Quantstudio 7 Flex using a 386-well plate (ThermoFisher Cat# 4,309,849) with Standard Mode. Target gene expression was normalized by the ΔΔCT method, and 18S was used as the internal control. qPCR primers used:

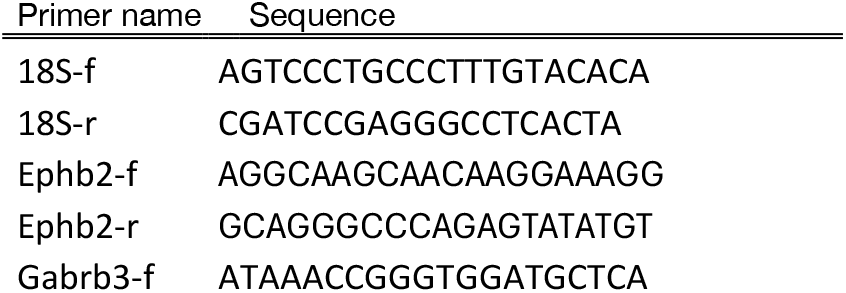
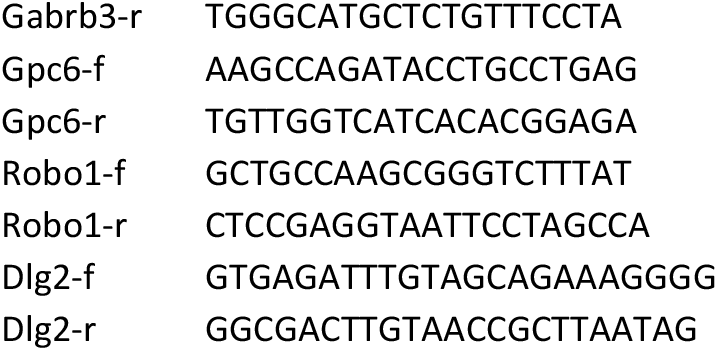

### Protein extraction and Western blotting

Total protein was extracted using 1xLDS loading buffer (1x NuPAGE LDS Sample Buffer (Invitrogen Cat# NP0007), 0.5% β-mercaptoethanol, 1 mM PMSF, EDTA-free Protease Inhibitor Cocktail (PI78439). Protein lysates were boiled at 70°C for 10 minutes and stored at -20 °C till used. Samples were pre-heated at 37 °C for 5 minutes and loaded to 4–12% NuPAGE™ Bis-Tris Mini Protein Gels (Invitrogen Cat# NP0322PK2) set in XCell SureLock Mini-Cell (Invitrogen Cat# EI0001) filled with 1x NuPAGE MOPS SDS Running Buffer (Invitrogen Cat# NP000102) and the electrophoresis was done using PowerPac Basic Power Supply (Bio-Rad Cat# 1645050). Proteins in the gel were transferred onto 0.45μm Immobilon-FL transfer membranes (Millipore Cat# IPFL00010) using an XCell II Blot Module (Invitrogen Cat# EI9051) filled with 1x NuPAG Transfer Buffer (Invitrogen Cat# NP00061) with 0.01% SDS and 5% methanol. After membranes were blocked with Intercept (PBS) Blocking Buffer (LI-COR Cat# 927-70001), proteins of interest were probed with specific primary antibodies and then appropriate Fluorescent Dye-conjugated secondary antibodies diluted in Intercept T20 (TBS) Antibody Diluent (LI-COR Cat# 927-75001). Fluorescent signals were detected using a ChemiDoc MP Imaging System (Bio-Rad Cat# 12003154). The following primary antibodies were used for immunoblotting: Upf2 (Cell Signaling Technology, Cat#11875), β-Actin (Cell Signaling Cat# 8457), phospho-Upf1 (Ser1127) antibody (MilliporeSigma, 07-1016), The following secondary antibodies were used: IRDye 800CW Goat anti-Rabbit IgG Secondary Antibody (LI-COR Cat# 926-32211) and IRDye 800CW Goat anti-Mouse IgG Secondary Antibody (LI-COR Cat# 926-32210)

## FIGURE LEGEND

**Supplementary Table 1**. The list of genes with intron/exon level differential expression at short/long-term HFD.

**Supplementary Table 2**. GO term analysis for intron/exon level DEGs at short-term HFD.

**Supplementary Table 3**. GO term analysis for intron/exon level DEGs at long-term HFD.

## Author Contributions

S.T. performed the bioinformatics analyses. M.J., S.T., performed the experiments and analyzed the data. S.T. and X.R. wrote the manuscript. X.R. conceived and supervised the study.

## Acknowledgments

We thank Dr. Bo Torben Porse for providing the Upf2^flox/flox^ mice. We are grateful to Dr. Hogg, J. Robert for helpful comments.

